# A *KCNC1* Variant Linked to Rett Syndrome Disrupts ER to Golgi Trafficking of Kv3.1 Channel

**DOI:** 10.1101/2024.11.12.623225

**Authors:** Diego Maureira, Carla Rubilar, Joaquín López, Paola Santander, Hans Moldenhauer, Ian Silva, Pablo Cruz, Denise Riquelme, Javiera Baeza, Wendy González, Patricio Orio, Evrim Servili, Mónica Troncoso, Elías Leiva-Salcedo, Oscar Cerda

## Abstract

Intrinsic neuronal excitability, defined by the balance between input and output signals, is crucial to neural function, and its disruption underlies various neurological diseases. Kv3.1 channels, encoded by *KCNC1*, are essential for high-frequency action potential firing. Variants in these channels are associated with several subtypes of epilepsy. We report a patient with developmental regression and epilepsy, meeting Rett syndrome criteria, who carries a *KCNC1* variant encoding the S474C substitution in Kv3.1 (Kv3.1^S474C^). Electrophysiological and biochemical assays reveal that Kv3.1^S474C^ reduces channel presence in the plasma membrane and is retained in the endoplasmic reticulum (ER). In murine primary cultures expressing Kv3.1^S474C^, we observed reduced neuronal firing frequency and exclusion of the channel from the axon initial segment (AIS). Consistently, we found a decreased firing frequency using a conductance-based computational neuronal model. In summary, this study identifies a novel link between a *KCNC1* variant and Rett syndrome, highlighting the importance of S474 residue in Kv3.1 channel trafficking and function in neurons.

**Summary:** This study identifies and characterizes a novel *KCNC1* variant associated with classical Rett syndrome. This variant disrupts endoplasmic reticulum (ER) to Golgi trafficking of the Kv3.1 channels, highlighting the variant’s potential role in altered neuronal excitability and neurodevelopmental disorders.

## Introduction

Potassium channels are encoded by more than 70 human genes, and dysregulation of these channels is implicated in numerous neurological, muscular, renal, and cardiac diseases (Lehmann-Horn & Jurkat-Rott, 1999). The *KCNC1* gene encodes two isoforms of the Kv3.1 channel, Kv3.1a and Kv3.1b, through alternative splicing (Liu & Kaczmarek, 1998). Variants in *KCNC1* have only recently been associated with pathological conditions, initially identified in a novel subtype of progressive myoclonic epilepsy termed “myoclonus epilepsy and ataxia due to potassium channel mutation” (MEAK) (Muona et al., 2015; Nascimento & Andrade, 2016; Oliver et al., 2017). Subsequently, *KCNC1* variants have been associated with other disorders, including developmental and epileptic encephalopathy, epilepsy of infancy with migrating focal seizures, and developmental delay without epilepsy (Ambrosino et al., 2023; Cameron et al., 2019; Li et al., 2021; Nascimento & Andrade, 2016; Park et al., 2019; Poirier et al., 2017). However, many *KCNC1* variants remain classified as variants of uncertain significance, requiring further clinical and genetic characterization to improve diagnostic accuracy and inform treatment strategies (Courage et al., 2021).

Kv3.1 is predominantly expressed in fast-firing GABAergic interneurons within the central nervous system (Gan & Kaczmarek, 1998). The channel’s biophysical properties, including a high activation threshold, rapid activation, and fast deactivation, facilitate swift repolarization during the action potential (AP) thereby enabling repetitive firing with brief AP durations. (Kaczmarek & Zhang, 2017; Pongs, 2008; Rudy & McBain, 2001). The loss of Kv3.1 function impairs the GABAergic interneurons firing, which may result in network hyperexcitability, a possible mechanism underlying various neurological disorders (Gunthorpe, 2022). In this report, we characterize a female patient with a heterozygous missense variant in *KCNC1*, presenting with developmental regression, stereotypic movements, congenital microcephaly, and epilepsy, consistent with classic Rett syndrome criteria. The identified variant, *KCNC1*: c.1421C>G, encodes a Ser474Cys substitution in the C-terminal region of the Kv3.1 that regulates the channel folding and trafficking. To date, no *KCNC1* variants have been reported to disrupt membrane trafficking and produce a pro-epileptic neuronal phenotype. This study demonstrates that the missense variant affecting the S474 residue in a patient meeting classical Rett syndrome criteria impairs normal anterograde trafficking of the channel, reduces Kv3.1 current, broadens the action potential, and reduces neuronal firing frequency.

## Results

### Clinical case characterization and sequence analysis results

The proband is a female, the fourth child of unrelated Chilean parents, born following a late-controlled pregnancy, with her first medical checkup occurring at an estimated gestational age of six months. Her father passed away from an acute myocardial infarction when she was 17 years old. Her mother, aged 38 at the time of delivery, had a history of one spontaneous abortion, substance use during pregnancy (including alcohol, tobacco, and benzodiazepines), recurrent preterm labor symptoms, gestational diabetes, and intrauterine growth restriction of unspecified cause. The patient was born at 36 weeks post-conception, weighing 2,000 g (below the 3rd percentile for the Chilean population), with a birth length of 43 cm (small for gestational age) and a head circumference of 29.8 cm (more than 2 standard deviations below the mean). Her Apgar score is unknown, and neither infections nor other significant perinatal complications were reported.

The patient demonstrated normal psychomotor development until 18 months of age when developmental stagnation became apparent. At 21 months, concurrent with an acute episode of pyelonephritis and distal renal tubular acidosis, she exhibited signs of developmental regression. These signs included loss of previously acquired language skills and social smiling, along with the emergence of hand flapping, stereotyped midline hand movements, and progressive gait deterioration. Her siblings are healthy, with no history of epilepsy, abnormal movements, tremors, or intellectual and developmental issues.

At two years of age, the patient experienced her first tonic-clonic seizures during a febrile episode. Subsequent seizures occurred without identifiable triggers, lasting approximately 10 minutes and occurring one to three times per day. Initially, seizure frequency was significantly reduced with valproic acid. At age seven, she was referred to our center for an etiologic evaluation. By that time, her phenotype had evolved to include severe intellectual disability, acquired microcephaly, synophrys, tapered fingers, short stature, scoliosis, and recurrent urinary tract infections. Based on her clinical history and behavioral profile, Rett syndrome was suspected.

The study began with a brain computed tomography (CT), which revealed diffuse brain atrophy with no other abnormalities (Figure 1A). Tandem mass spectrometry of blood showed only an elevated methionine concentration. An electroencephalogram (EEG) showed poor representation of physiological sleep elements without epileptiform abnormalities. Polysomnography was normal. Genetic evaluation included a normal karyotype and Sanger sequencing of the *MECP2* gene, which revealed no mutations but did identify the c.1233C>T (p.Pro411=) synonymous variant. Additional tests, including FISH for subtelomeric rearrangements, MS-MLPA for *CDKL5*, *NTNG1*, *ARX*, and *MECP2*, as well as an Angelman Syndrome and Related Disorders NGS Panel (Invitae Corporation) were all normal.

**Figure 1.**
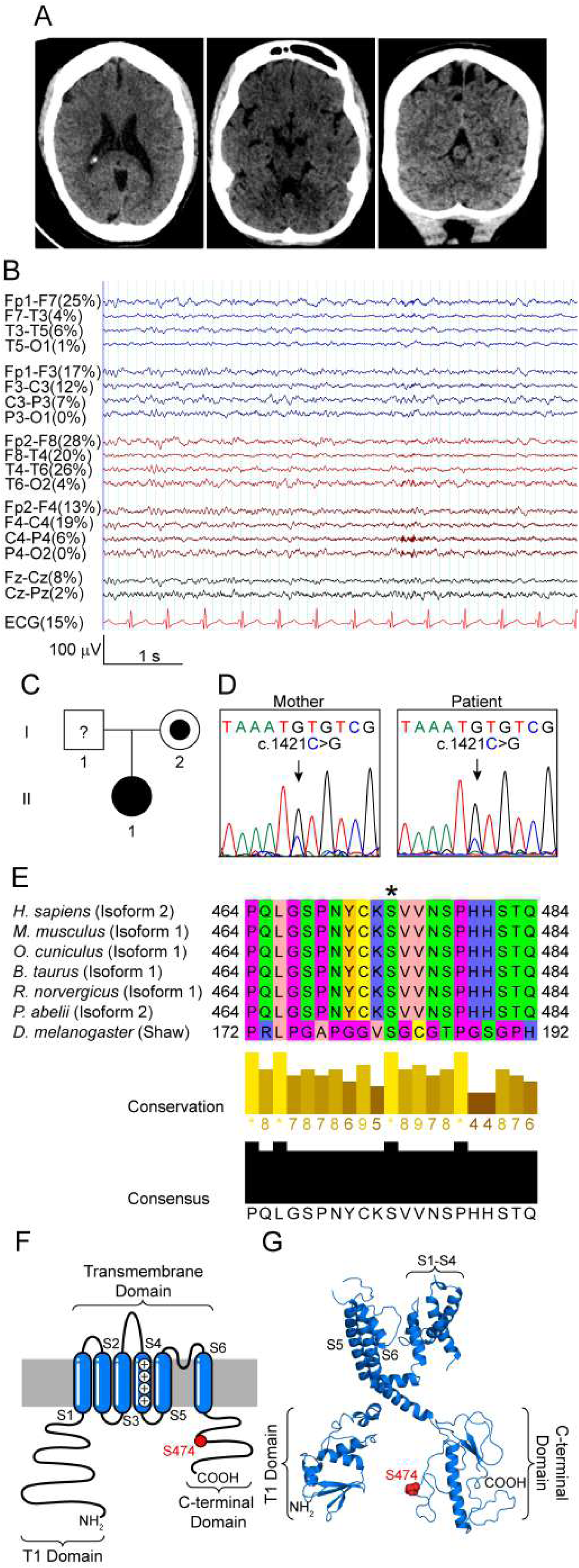
Kv3.1b^S474C^ variant found in a patient with classical Rett syndrome criteria. **(A)** and **(B)** Brain CT and EEG obtained from the patient, where diffuse cortex atrophy and no epileptogenic abnormalities was observed respectively. **(C)** Pedigree of Patient with Kv3.1 channelopathy. Filled circles (females) and squares (Citri & Malenka) indicate heterozygosity; symbols with “?” indicate an asymptomatic member without available DNA sequencing. **(D)** Electropherogram of genomic DNA obtained from the mother and the proband. **(E)** Sequence alignment of Kv3.1 protein sequences from *Homo sapiens* (human), *Mus musculus* (mouse), *Oryctolagus cuniculus* (rabbit), *Bos taurus* (cattle), *Rattus norvergicus* (rat) *Pongo abelii* (orangutan) and *Drosophila melanogaster* (fly). A multiple-sequence alignment was built with ClustalW server and analyzed with Jalview v.1.8.3. The sequence alignment is colored according to the chemical properties of the residues. Conservation of the S474 residue is indicated (asterisk). Conservation and consensus are indicated in the bottom diagrams **(F)** Membrane topology of the Kv3.1b channel. The plasma membrane is presented in gray and the residue S474 in red. **(G)** Structural topology of the Kv3.1b channel is shown as ribbons. Residue S474 is presented as sphere model in red.

The patient exhibited progressive loss of purposeful hand skills, autistic behaviors, inappropriate laughing, and self- and hetero- aggression, which were partially controlled with risperidone. She also demonstrated loss of spoken language, profound intellectual disability, intense eye contact, and lower limb spasticity. At six years of age, she lost her ability to walk but later regained partial gait function, with a rigid gait, an increased base of support, and a forward-shifted center of gravity. Other clinical features included malnutrition, short stature, scoliosis, bruxism while awake, and recurrent urinary tract infections. Seizures were well-controlled with dose adjustments of valproic acid monotherapy, with tonic-clonic episodes occurring only during febrile states unrelated to epilepsy. EEGs remained stable, showing no epileptiform activity, and continued poor representation of physiological sleep elements (Figure 1B).

Given the negative results of previous studies, an epilepsy Next Generation Sequencing (NGS) panel (Invitae Corporation) was conducted, identifying two heterozygous variants of uncertain significance: *KCNC1* c.1421C>G (p.Ser474Cys) and *RELN* c.7648A>G (p.Met2550Val) (Figure 1C-D). Subsequent sequencing of the proband’s mother revealed that she was also a heterozygous carrier of the *KCNC1* c.1421C>G variant, although she had no history of epilepsy (Figure 1C-D). Further segregation analysis could not be performed because the proband’s father was deceased. The classification of these variants, using the American College of Medical Genetics and Genomics criteria (ACMG) (Richards et al., 2015) was carried out via the Varsome database (Kopanos et al., 2019) using Human Phenotype Ontology IDs: HP:0002069, HP:0002376, HP:0004322, HP:0000733, HP:0000718, HP:0002650, HP:0000664, HP:0001238, and HP:0000010 (The Human Phenotype Ontology (Brittain et al., 2011) https://hpo.jax.org/app/; (Köhler et al., 2019)). Both variants were classified as variants of uncertain significance (VUS).

The classification of the *KCNC1* variant was based on its allele count of one in population databases and its status as a missense variant in a gene with a low frequency of benign variants. In contrast, the *RELN* variant had an allele count of three in the GnomAD 4.0 database (Chen et al., 2024), and another variant affecting the same amino acid (p.Met2550Thr) was identified in the same database with an allele count of one. A manual sequence alignment analysis indicated that residue S474 of Kv3.1 is highly conserved among species, whereas the Met2550 residue of Reelin is located in a more variable region (Figure 1E, Supplementary Figure 1A). For the *RELN* variant, *in silico* predictors from the dbNSFP v4.8 database (Liu et al., 2011; Liu et al., 2020) accessed *via* Varsome, returned unanimous benign meta-scores. Conversely, for the *KCNC1* variant, the meta-scores returned were classified as “pathogenic” (BayesDel addAF, MetaLR, and BayesDel noAF) or “uncertain” (MetaRNN, MetaSVM, and REVEL) (Supplementary Figure 1B).

The p.Ser474Cys variant impacts in the C-terminal of the channel, as was evidenced by a Kv3.1b topology model (Figure 1F-G), which is crucial for its trafficking and targeting within neurons (Xu et al., 2007). Given the clinical context and *in silico* prediction data, we decided to investigate the impact of this *KCNC1* variant on Kv3.1 function and its effect on neuronal physiology.

### Kv3.1b^S474C^ reduces the current density of the Kv3.1b channel

Kv3.1 currents are crucial in regulating neuronal excitability by contributing to the rapid repolarization of action potentials, shortening spike duration, and enabling high-frequency firing. Kv3.1 is a delayed rectifier K^+^ channel characterized by a minimal relative refractory period (Grissmer et al., 1994; Kaczmarek & Zhang, 2017; Kanemasa et al., 1995; Luneau et al., 1991). To assess the effect of the *KCNC1*: c.1421C>G (p.Ser474Cys) variant in Kv3.1b activity, we expressed FLAG-Kv3.1b wild-type (FLAG-Kv3.1b^WT^) or the patient-identified variant (FLAG-Kv3.1b^S474C^) in HEK293 cells. Whole-cell patch-clamp recordings from cells expressing FLAG-Kv3.1b^S474C^ showed a 60% reduction in current density compared to wild-type, with mean current values (at 40mV) of 98 ± 29 pA/pF (n=18 cells) for FLAG-Kv3.1b^WT^ and 45 ± 16 pA/pF (n=16 cells) for FLAG-Kv3.1b^S474C^ (Figure 2A-B). No significant differences in the half-maximal activation voltages (V_0.5_) were observed (+12 ± 15 mV, n=18 cells for FLAG-Kv3.1b^WT^ and +20 ± 17 mV, n=16 cells for FLAG-Kv3.1b^S474C^), indicating that the reduced current density of the *KCNC1*: c.1421C>G (p.Ser474Cys) variant is not due to changes in the voltage dependence of the channel (Figure 2C-D).

**Figure 2.**
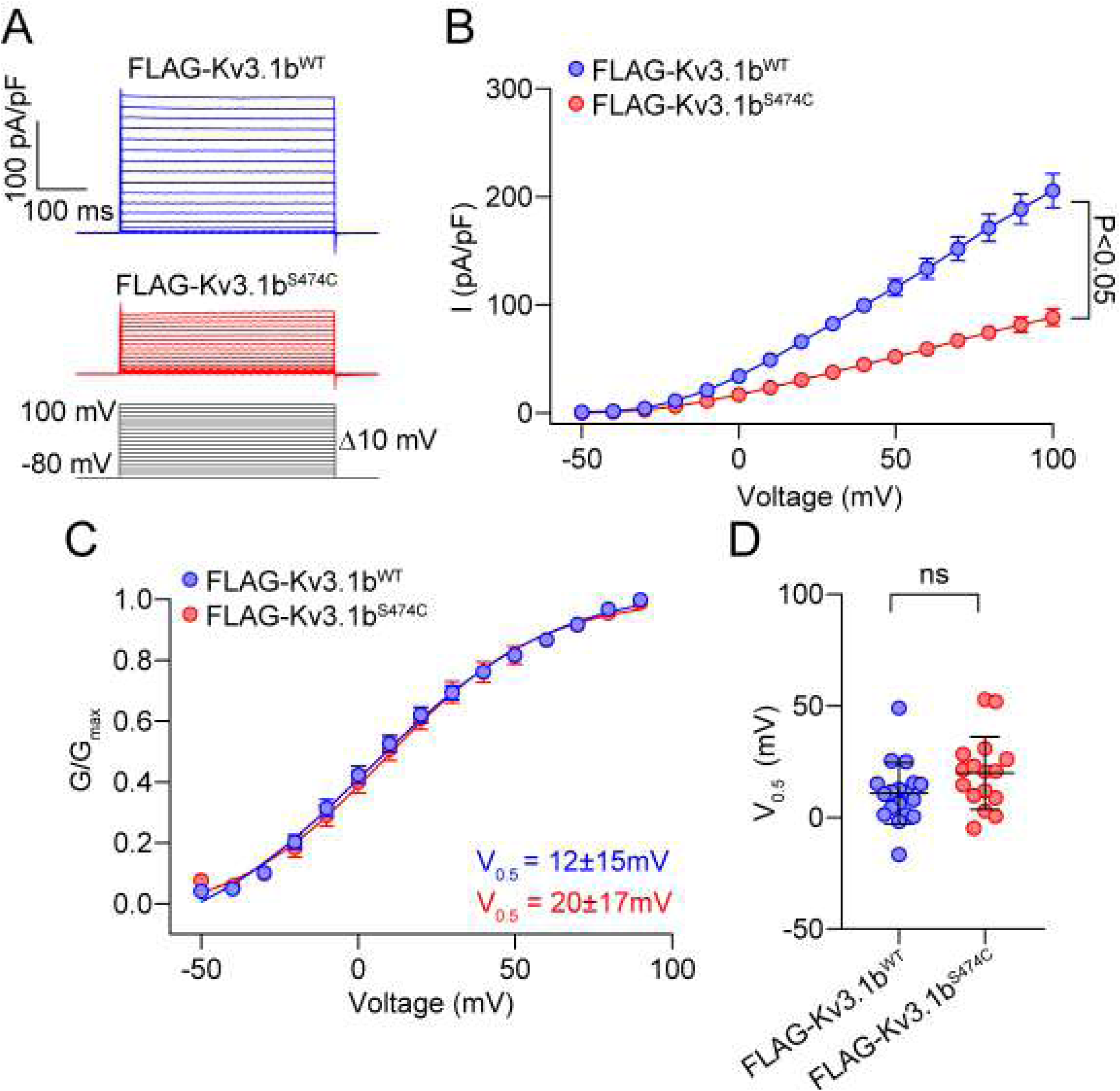
Kv3.1b^S474C^ substitution reduces the Kv3.1b currents. **(A)** Representative current traces from HEK293 cells expressing FLAG-Kv3.1b^WT^ and FLAG-Kv3.1b^S474C^, evoked by voltage steps from −80 mV to +100 mV with a 10 mV increments, from a holding potential of −80 mV. **(B)** Average current-voltage (I/V) curves (mean±SEM) of FLAG-Kv3.1b^WT^ and FLAG-Kv3.1b^S474C^ expressed in HEK293 cells, significant differences were observed since −20 mV. Statistical analysis was performed using two-way ANOVA, followed by Tukey’s post hoc test. **(C)** Conductance-voltage (G-V) relationship for FLAG-Kv3.1b^WT^ (n=18 cells) and FLAG-Kv3.1b^S474C^ (n=16 cells) channels. G was normalized to the highest conductance calculated (G_max_). The G-V curves were fitted by a Boltzmann function. **(D)** Half-maximal activation potential (V_0,5_) values for FLAG-Kv3.1b^WT^ and FLAG-Kv3.1b^S474C^channels. Statistical analysis was performed using a nonparametric Mann-Whitney test. n.s., not significant.

### Kv3.1b^S474C^ exhibits a reduced number of channels in the plasma membrane

Current density is determined by several factors, including the number of channels, unitary current, and open probability (Hille, 2001). Noise analysis, a method that allows for the estimation of single-channel activity, was performed to assess the plasma membrane population of ion channels (Alvarez et al., 2002). These analyses revealed a significant reduction in the number of channels in the plasma membrane (139,000 ± 34,000 channels, n=10 cells for FLAG-Kv3.1b^WT^ and 69,000 ± 23,000 channels, n=9 cells for FLAG-Kv3.1b^S474C^) (Figure 3A-B). No changes were observed in the unitary current, with values (63 ± 19 fA, n=10 cells for FLAG-Kv3.1b^WT^ and 71 ± 32 fA, n=9 cells for FLAG-Kv3.1b^S474C^) (Figure 3C). To corroborate these findings, we performed surface-biotinylation assays in HEK293 cells expressing either FLAG-Kv3.1b^WT^ or FLAG-Kv3.1b^S474C^. Consistent with the noise analysis, the biotinylated population of FLAG-Kv3.1b^S474C^ was reduced compared to FLAG-Kv3.1b^WT^, indicating a decreased plasma membrane expression of the mutant channel (Figure 3D-E).

**Figure 3.**
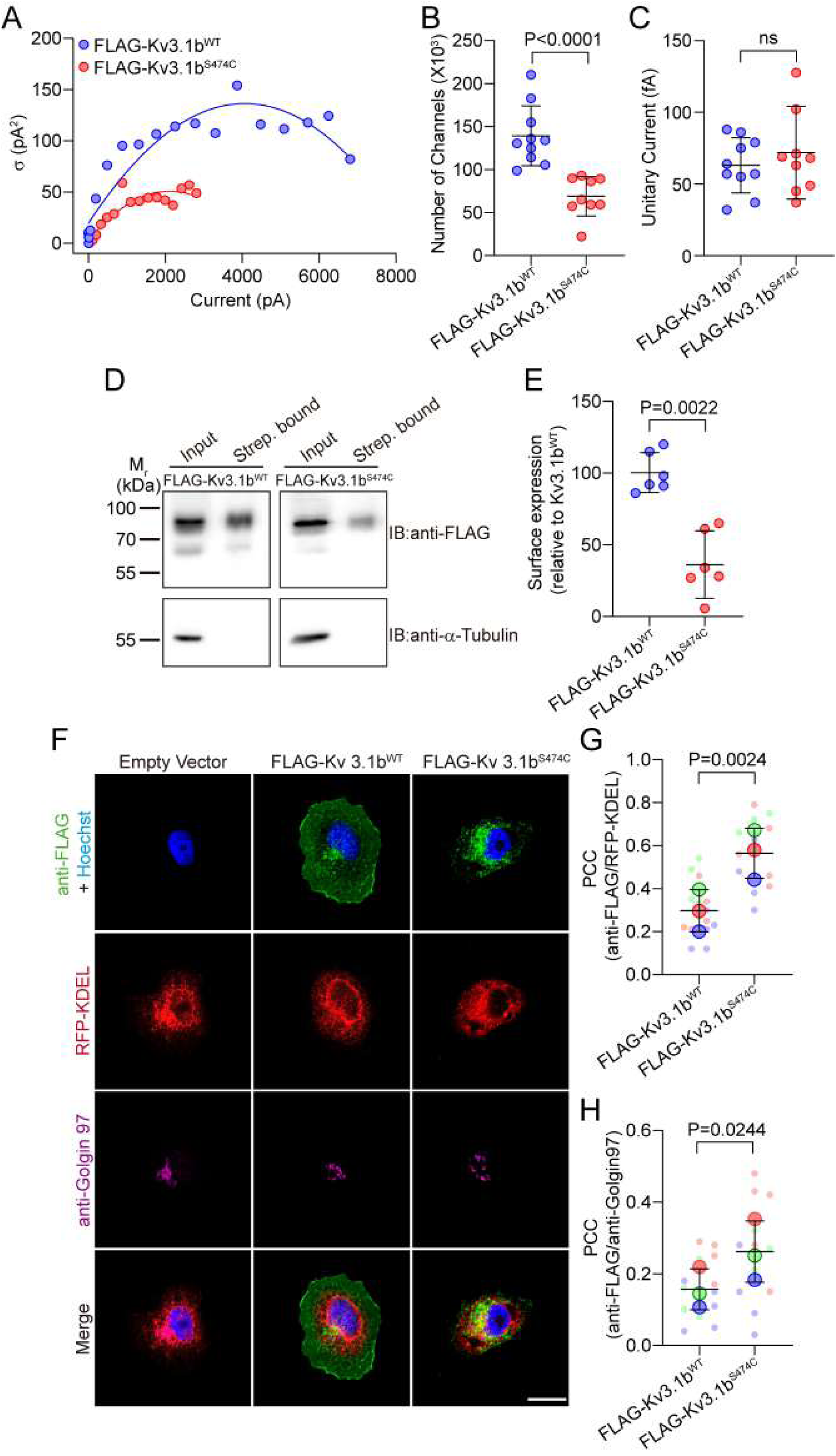
Kv3.1b^S474C^ substitution in Kv3.1b reduces the number of channels in the plasma membrane. **(A)** Representative traces of non-stationary noise analysis are shown for FLAG-Kv3.1b^WT^ and FLAG-Kv3.1b^S474C^ channels, where mean current (pA) is plotted against current variance σ (pA^2^). Curves were fitted with a second-order polynomial function. **(B)** Number of channels and **(C)** unitary currents for FLAG-Kv3.1b^WT^ (n=10 cells) and FLAG-Kv3.1b^S474C^ (n=9 cells) channels obtained from noise analysis. Statistical analysis was performed using the nonparametric Mann-Whitney test. n.s., not significant. **(D)** Cell-surface biotinylation assays from HEK293 cells expressing FLAG-Kv3.1b^WT^ and FLAG-Kv3.1b^S474C^ constructs. A representative image of results obtained from 6 independent experiments is shown. Membranes were incubated with anti-FLAG, anti-TRPM4, and anti-α-tubulin antibodies. **(E)** The graph shows the surface expression (mean±SD) signal intensity of Kv3.1b levels in the streptavidin-bound fraction relative to the input and normalized against FLAG-Kv3.1b^WT^. Statistical analysis was performed using the nonparametric Mann-Whitney test. **(F)** Representative images of HeLa cells cotransfected with RFP-KDEL (red) as an ER marker and either FLAG-Kv3.1b^WT^ or FLAG-Kv3.1b^S474C^ constructs. The cells were immunolabeled for the Kv3.1b FLAG tag (green), the Golgi apparatus with anti-golgin 97 (magenta), and the nuclei with Hoechst 33342 (blue). **(G)** Pearson’s correlation coefficient (PCC) (mean±SD) of ER colocalization with FLAG-Kv3.1b^WT^ (n= 16 cells) and FLAG-Kv3.1b^S474C^ (n=15 cells) constructs. **(H)** Pearson’s correlation coefficient (mean±SD) of Golgi colocalization with FLAG-Kv3.1b^WT^ (n=15 cells) and FLAG-Kv3.1b^S474C^ (n=15 cells) constructs. Statistical analysis was performed using the nonparametric Mann-Whitney test.

Since the Kv3.1b^S474C^ substitution of Kv3.1b leads to a reduction in the number of channels in the plasma membrane, we further investigated the subcellular localization of these channels. Immunofluorescence assays were performed in HeLa cells overexpressing FLAG-Kv3.1b^WT^ or FLAG-Kv3.1b^S474C^, along with the ER marker RFP-KDEL and the Golgi apparatus marker anti-golgin 97 (Figure 3F). We observed an increase in the Pearson Correlation Coefficient (PCC) between FLAG-Kv3.1b^S474C^, and both ER and Golgi markers (Figure 3G-H), suggesting that Kv3.1b^S474C^ is retained intracellularly, specifically in the ER and Golgi apparatus.

### Kv3.1b^S474C^ impairs Kv3.1b channel trafficking beyond the ER

To gain further insight into the role of the *KCNC1*: c.1421C>G (p.Ser474Cys) variant in Kv3.1b trafficking, we synchronized the anterograde trafficking of the channel using 20 μM Nocodazole (Storrie, 1998), which inhibits the polymerization of microtubules, in HeLa cells transfected with FLAG-Kv3.1^WT^ or FLAG-Kv3.1^S474C^ and the RFP-KDEL ER marker. After 16 hours of synchronization, we washed out the drug from the media and the cells were fixed at different times. Interestingly, after 60 minutes of the remotion of the drug, we observed a decrease in the PCC between FLAG-Kv3.1^WT^ and the RFP-KDEL marker, while the FLAG-Kv3.1^S474C^ retained its colocalization with RFP-KDEL after 60 minutes of washout (Figure 4A-B). We also used WGA conjugated to Alexa fluor 555 to stain the plasma membrane in synchronized Hela cells transfected with FLAG-Kv3.1^WT^ or FLAG-Kv3.1^S474C^. After 60 minutes of Nocodazole washout, we observed an increase in the PCC between the FLAG-Kv3.1^WT^ and the membrane marker, while FLAG-Kv3.1^S474C^ did not increase (Figure 4C-D). Moreover, we used a different approach to synchronize the anterograde trafficking of the channel. As such, we used the Retention Using Selective Hooks (RUSH) system in HeLa cells transfected with either Kv3.1b^WT^-mVenus-SBP or Kv3.1b^S474C^-mVenus-SBP along with the bicistronic Str-KDEL-IRES-mScarlet-i-Sec23a construct, which encodes an ER-associated Hook and the ER exit site (ERES) marker mScarlet-i-Sec23a (Boncompain et al., 2012; McCaughey et al., 2019). After D-biotin incubation, the cells were fixed and stained with an anti-Golgin 97 antibody as a Golgi marker (Figure 4A). Notably, we observed an increase in the PCC between the wild-type Kv3.1b and the ERES marker 30 minutes after D-biotin addition. After 60 minutes, there was an increase in the PCC between Kv3.1b^WT^-mVenus-SBP and Golgin 97 marker (Figure 4D-E). In contrast, no increase in the PCC was observed between Kv3.1b^S474C^-mVenus-SBP, and either Sec23 or Golgin 97 (Figure 4F-G), indicating that the S474 residue is critical for the channel’s entry into the ERES and, consequently, its anterograde trafficking.

**Figure 4.**
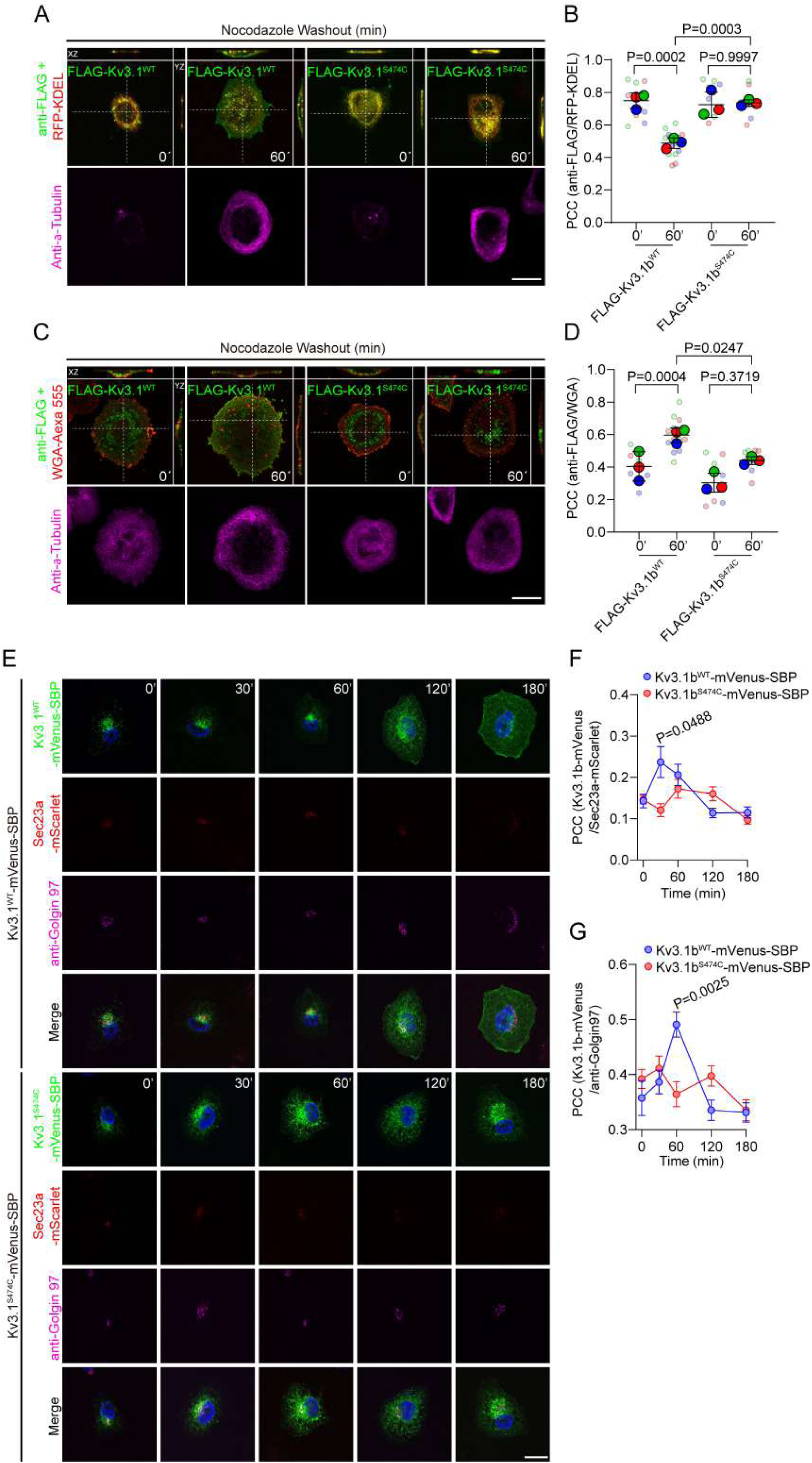
Kv3.1b^S474C^ substitution in Kv3.1b retains the channels in the ER. **(A)** Representative images of HeLa cells coexpressing RFP-KDEL (red) and FLAG-Kv3.1b^WT^ and FLAG-Kv3.1^S474C^ before (0 min) and after (60 min) nocodazole-washout (20 µM), with orthogonal views from different planes wqdds’’’’’x/y, x/z, or y/z). The cells were immunolabeled with anti-FLAG, (green), anti-α-tubulin (magenta), and Hoechst 33342 (blue). **(B)** Pearson’s correlation coefficient (PCC) (mean±SD) of ER colocalization with FLAG-Kv3.1^WT^ at 0 min (n=10 cells) or 60 min (n=15 cells), and with FLAG-Kv3.1^S474C^ at 0 min (n=10 cells) and 60 min (n=11 cells), respectively. **(C)** Representative images of HeLa cells expressing Kv3.1b^WT^ or Kv3.1^S474C^ before (0 min) and after (60 min) nocodazole-washout, with orthogonal views from different planes (x/y, x/z, or y/z). The cells were immunolabeled for the Kv3.1b FLAG tag (green), the tubulin cytoskeleton (magenta), the membrane with WGA (red), and the nuclei with Hoechst 33342 (blue). **(D)** PCC (mean±SD) of the membrane (WGA) colocalization with FLAG-Kv3.1^WT^ at 0 min (n=10 cells) or 60 min (n=13 cells), and with FLAG-Kv3.1^S474C^ at 0 min (n=10 cells) and 60 min (n=14 cells), respectively. Statistical analysis was performed using ANOVA followed by Dunnett’s multiple comparison test **(E)** Representative images of HeLa cells cotransfected with Str-KDEL-IRES-mScarlet-i-Sec23a (red) as a ERES marker and Kv3.1b^WT^-mVenus-SBP or Kv3.1b^S474C^-mVenus-SBP. The cells were immunolabeled with anti-Golgin 97 (magenta) as a Golgi marker. **(F)** PCC (mean±SEM) of mScarlet-i-Sec23a colocalization with Kv3.1b^WT^-mVenus-SBP (n=73 cells) or Kv3.1b^S474C^-mVenus-SBP (n=76 cells) constructs. Statistical analysis was performed using two-way ANOVA, followed by Tukey’s post hoc test. **(G)** PCC (mean±SEM) of Golgi colocalization with Kv3.1b^WT^-mVenus-SBP (n=73 cells) or Kv3.1b^S474C^-mVenus-SBP (n=76 cells) constructs. Statistical analysis was performed using two-way ANOVA, followed by Tukey’s post hoc test.

### Kv3.1b^S474C^ alters the ion channel folding

Next, we investigated whether the S474 residue participates in ion channel folding. Although no crystallographic structure of Kv3.1b is available, we generated a Kv3.1b channel model using homology modeling. Kv3.1a and Kv3.1b channels share 98.24% sequence identity; the difference resides in the C-terminal segments beginning at residue 502. Therefore, the canonical structure of the Kv3.1 channel (residues 1-454; PDB ID: 7PQT) was used as a template, and the missing C-terminal fragment corresponding to the Kv3.1b channel (*i.e.*, residues 454-549) was generated using CollabFold. After reaching the thermodynamic equilibrium, a 500 ns molecular dynamics simulation was performed for Kv3.1b^WT^, where the Root Mean Standard Deviation (RMSD) remained stable at 8∼8.5 Å. We analyzed the secondary structure of the constructed model and observed that the structure of the Kv3.1b channel has segments similar to the consensus secondary structure prediction (Supplementary Figure 2). Although some differences were observed, probably due to local changes contributed by the molecular dynamics simulation, we identified secondary structures that suggest a similar folding pattern. Comparing the secondary structure prediction from molecular dynamics-based structure with the sequence-based secondary structure prediction, we found consistent patterns. A similar simulation was launched for the Kv3.1b^S474C^ substitution using the same configuration. Unlike the wild-type system, the Kv3.1b^S474C^ substitution did not reach a plateau and appeared to approach equilibrium only at the final stages of the simulation (Figure 5A). Also, our molecular dynamics showed that the Kv3.1b^S474C^ substitution delays the folding of the C-terminal segment compared to Kv3.1^wt^ (Figure 5B). Importantly, an increased number of contacts were obtained in the Kv3.1b^WT^ system compared to the Kv3.1b^S474C^ system (*i.e.*, 39 versus 31 interactions found for the Kv3.1b^WT^ and Kv3.1b^S474C^ systems, respectively). Furthermore, in the Kv3.1b^WT^ system, the S474 residue interacts with the N-terminal domain of the adjacent subunit (Figure 5C-D). In contrast, in the Kv3.1b^S474C^ substitution, residue C474 maintains interactions mainly with the surrounding intramolecular residues (Figure 5C). No previous reports of specific interactions were identified for the described residues. Additionally, to further explore whether the *KCNC1*: c.1421C>G (p.Ser474Cys) variant affects the free energy and folding stability of the native channel, we conducted computational analyses using several single mutation prediction servers. These analyses indicated that the Kv3.1b^S474C^ substitution destabilizes channel folding (Figure 5D).

**Figure 5.**
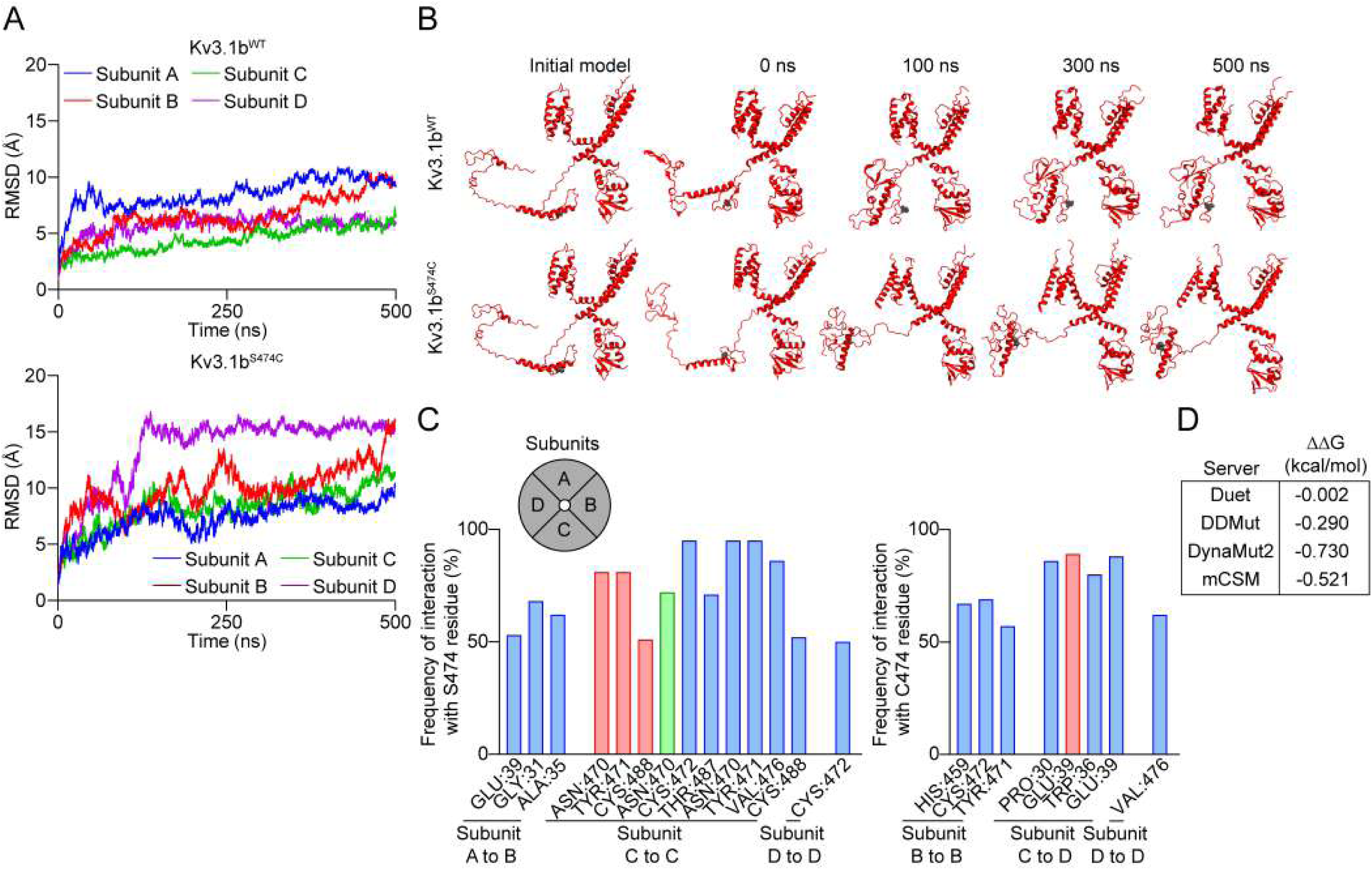
Kv3.1b^S474C^ substitution in Kv3.1b affects the channel folding. **(A)** RMSD of the backbone from Kv3.1b^WT^ and Kv3.1b^S474C^ structures in molecular dynamics, respectively. **(B)** Snapshots of different frames along the molecular dynamics for the systems Kv3.1b^WT^ and Kv3.1b^S474C^ in cartoon representation. Residue 474 is shown in gray. **(C)** Analysis of intermolecular interactions, GetContacts program was used to extract the key amino acids for intermolecular interactions in the molecular dynamics simulations of systems Kv3.1b^WT^ and Kv3.1b^S474C^. The residues are shown in licorice representation. Residue 474 is shown in gray. Red and blue residues belong to chains A and B, respectively. **(D)** Main interactors residues observed in the 500 ns simulation in the S474 of Kv3.1b^WT^ or the C474 of Kv3.1b^S474C^. The red columns represent hydrogen bonds with the sidebone, green for hydrogen bonds with the backbone and blue for Van der Waals interactions with the respective residue. **(E)** Effect of the S474C mutation on the KV3.1b channel structure model. The difference in binding energy (ΔΔG) was calculated by performing the S474C mutation on DUET, DDMut, DynaMut2 and mCSM single-point mutation servers.

### Reduced firing frequency in neurons expressing the Kv3.1b^S474C^

Given that the axon initial segment (AIS) plays a critical role in regulating AP frequency (Yamada & Kuba, 2016) and that Kv3.1b is expressed in the AIS, promoting neuronal firing (Gu et al., 2012; Xu et al., 2007; Xu et al., 2010), we evaluated the role of the S474 residue in Kv3.1b trafficking in neurons. Immunofluorescence assays were performed in cortical primary cultures from mouse embryos transfected with FLAG-Kv3.1b^WT^ or FLAG-Kv3.1b^S474C^ and using anti-Ankyrin-G antibody as an AIS marker (Figure 6A). Notably, FLAG-Kv3.1b^S474C^ was excluded from the axonal domain, unlike the wild-type channel. This was demonstrated by a decreased PCC between the ion channel signal and the AIS marker (Figure 6B), indicating that S474 is essential for targeting Kv3.1b to the AIS. Importantly, no changes in AIS morphology was observed (Figure 6C).

**Figure 6.**
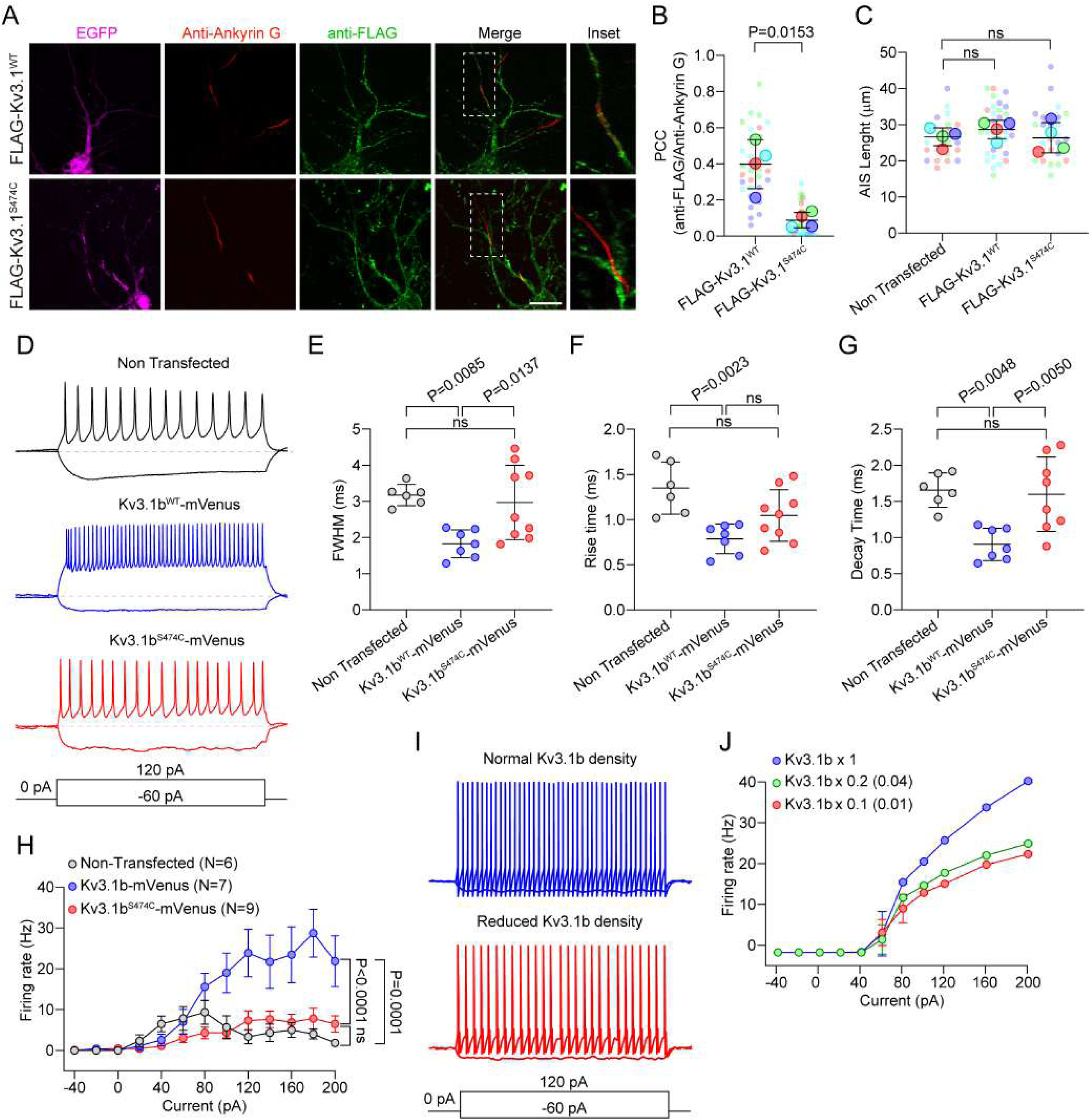
Kv3.1b^S474C^ substitution in Kv3.1b reduces the action potential firing. **(A)** Representative images of primary cortical neurons in culture transfected with FLAG-Kv3.1b^WT^ or FLAG-Kv3.1b^S474C^ constructs and EGFP. The cells were immunolabeled for the Kv3.1b FLAG tag (green), the AIS with anti-Ankyrin G (red), and the nuclei with Hoechst 33342 (blue). **(B)** Pearson’s correlation coefficient (PCC) (mean±SD) of AIS colocalization with FLAG-Kv3.1b^WT^ (n=32 cells) and FLAG-Kv3.1b^S474C^ (n=31 cells) constructs. Statistical analysis was performed using the nonparametric Mann-Whitney test. **(C)** AIS length (mean±SD) of neurons non-transfected (n=20) or transfected with FLAG-Kv3.1b^WT^ (n=33 cells) or FLAG-Kv3.1b^S474C^ (n=28 cells) constructs. Statistical analysis was performed using ANOVA followed by Dunnett’s multiple comparison test. n.s., not significant. **(D)** Representative traces of cortex primary culture neurons transfected with Kv3.1b^WT^-mVenus or Kv3.1b^S474C^-mVenus constructs evoked by current steps of −60 pA and +120 pA. The dotted line indicates the resting potential of any neuron (−63 mV for non-transfected, −64 mV for Kv3.1b^WT^-mVenus and −61 mV for Kv3.1b^S474C^-mVenus). **(E)** Action potential duration expressed as a Full width at half maximum (FWHM), **(F)** rise time, and **(G)** decay time of neurons non-transfected (n=6) or transfected with Kv3.1b^WT^-mVenus (n=7 cells) or Kv3.1b^S474C^-mVenus (n=9 cells). Statistical analysis was performed using ANOVA followed by Dunnett’s multiple comparison test. n.s., not significant. **(H)** Firing frequency (mean±SEM) of neurons non-transfected (n=6) or transfected with Kv3.1b^WT^-mVenus (n=7 cells) or Kv3.1b^S474C^-mVenus (n=9 cells). Statistical analysis was performed using two-way ANOVA, followed by Tukey’s post hoc test. **(I)** Simulated voltage traces in the presence of normal (0.2 / 0.8 mS/cm^2^; somatodendritic/axonal) or reduced (0.02 / 0.008 mS/cm^2^) Kv3.1b density. **(J)** Firing frequency of simulated neurons containing different densities of Kv3.1. Basal density is 0.016 and 0.064 mS/cm^2^ (somatodendritic and axonal, respectively). Number in the legend indicates the reduction factor for the density, first number for somatodendritic compartments and between parenthesis for the axonal compartment.

Kv3.1b has been identified as a critical component for high-frequency firing in neurons (Kaczmarek & Zhang, 2017; Pongs, 2008; Rudy & McBain, 2001). Overexpression of Kv3.1b in pyramidal neurons has been shown to increase firing frequency (Gu et al., 2018). In this context, we evaluated the role of S474 in AP firing through current-clamp recordings on cortical primary neuronal cultures from mouse embryos transfected with Kv3.1b^WT^-mVenus or Kv3.1b^S474C^-mVenus (Figure 6D). The expression of Kv3.1b^WT^-mVenus reduces the full-width half maximum (FWHM) (Figure 6E), rise time (Figure 6F), and decay time (Figure 6G), leading to an increase in firing rate (Figure 6H), consistent with previous studies (Kaczmarek & Zhang, 2017; Pongs, 2008; Rudy & McBain, 2001). In contrast, Kv3.1b^S474C^-mVenus showed values similar to those observed in non-transfected neurons (Figure 6D-H), indicating that the S474C variant is non-functional in neurons.

To further investigate the role of axonal Kv3.1b in enabling high-frequency firing, we performed conductance-based modeling of neuronal excitability. Based on the model developed by (Gu et al., 2012), we constructed a model neuron with dendritic, somatic, and axonal compartments, each containing a voltage-gated sodium (NaV) channel, a generic delayed rectifier potassium (Kdr) channel, and a voltage-gated potassium channel with Kv3.1b properties (Gu et al., 2012). The model was calibrated to reflect typical membrane resistance and capacitance values observed in cultured neurons, and simulations were used to test the dependence of firing rate on Kv3.1b density. A decrease in firing frequency was observed when Kv3.1b density was preferentially reduced in the axon (Figure 6I-J), mimicking the reduction observed in neurons expressing the S474C variant (Figure 6H). This effect was not observed when Kv3.1b was reduced in the somatic and dendritic compartments (data not shown). Similarly, reducing Kdr current density shifted the firing threshold without affecting the maximum firing rate (data not shown).

## Discussion

In this work, we identified a novel mutation in the KCNC1 gene in a patient with clinically diagnosed Rett syndrome but without alterations in previously reported genes such as MECP2, CDKL5, and FOXG1. This c.1421C>G (p.Ser474Cys) mutation results in the retention of the channel in the endoplasmic reticulum and Golgi apparatus, thereby reducing Kv3.1 membrane expression and K^+^ current. This, in turn, decreases neuronal firing. To our knowledge, this is the first described mutation in Kv3.1 that leads to intracellular channel retention and is associated with a pathological phenotype.

In this study, we conducted a thorough clinical and molecular evaluation of a patient who met the diagnostic criteria for classic Rett syndrome. Despite presenting hallmark symptoms of Rett syndrome (Kaur et al., 2019), genetic testing for the common Rett syndrome-associated genes *MECP2*, *CDKL5*, and *FOXG1* yielded negative results. However, further genetic analysis revealed a novel variant in the *KCNC1* gene (c.1421C>G (p.Ser474Cys)). Although this *KCNC1* variant has not been previously linked to Rett syndrome, recent advances in NGS, bioinformatics tools, and whole exome sequencing (WES) have uncovered novel genes associated with the disorder. Notably, several ion channels, such as *SCN2A*, *KCNQ2*, *SCN8A*, *KCNB1*, and *CACNA1I*, have all been implicated in the pathophysiology of Rett syndrome (Vidal et al., 2019).

Several variations which cause loss of function in *KCNC1* have been classically related to progressive myoclonic epilepsy (Ambrosino et al., 2023; Cameron et al., 2019; Franceschetti et al., 2014; Muona et al., 2015). Additionally, other neurological disorders such as epilepsy, ataxia, and developmental delay have been related to *KCNC1* variants (Cameron et al., 2019; Li et al., 2021; Park et al., 2019; Poirier et al., 2017; Zhang et al., 2021), although most of these variants are predicted or functionally characterized as loss-of-function. Interestingly, Clatot et al. (Clatot et al., 2023) identified three de novo heterozygous *KCNC1* gain-of-function variants (p.Met430Ile, p.Val432Met, and p.Val434Leu) in patients with developmental delay and hypotonia but without epilepsy or myoclonus. Similarly, Ambrosino et al. (Ambrosino et al., 2023) described a patient with developmental and epileptic encephalopathy who experienced fever-triggered seizures (p.Val425Met). This variant also exhibited a gain-of-function phenotype, in contrast to the typical loss-of-function variants seen in most cases.

Based on the patient’s genetic sequencing results, we performed an ACMG criteria analysis for both the *KCNC1* and *RELN* variants using the Franklin by Genoox database (https://franklin.genoox.com/clinical-db/home). Both variants were classified as VUS, with the *KCNC1* variant potentially changing classification to likely pathogenic if further functional studies provide additional evidence. Notably, the proband’s unaffected mother is a carrier of the *KCNC1* variant, which is unusual since most reported pathogenic *KCNC1* variants are de novo. This raises at least three potential explanations. The first explanation regarding the mother’s unaffected status suggests that the outcome of the variant in the mother may depend on the genomic background, which could potentially rescue the *KCNC1*:c.1421C>G (p.Ser474Cys) variant through interacting proteins. This could modulate the phenotype and protect the mother from developing a pathological phenotype. The second explanation considers the combined effect of the *RELN* and *KCNC1* variant, although we did not assess its potential combined impact on neuronal function. We also did not assess the *RELN* variant carrier status in the proband’s mother and, therefore, we cannot exclude its contribution. The third explanation suggests that prenatal factors such as maternal substance abuse, small for gestational age status, and subsequent failure to thrive may have contributed to the patient’s pathological phenotype. Finally, we propose that the *KCNC1* variant could be the primary driver for several reasons: functional studies demonstrate that the variant leads to Kv3.1 loss-of-function in neuronal models, in silico predictions suggest the *RELN* variant is benign, and the proband’s clinical features align more closely with those observed in patients carrying *KCNC1* variants rather than *RELN*-associated conditions, such as Lissencephaly type 2 (OMIM #616436) or familial temporal lobe epilepsy-7 (OMIM #257320) (Di Donato et al., 2022). Based on these findings, we hypothesize that the patient’s clinical manifestations are more likely due to Kv3.1 dysfunction rather than a Reelin-dependent effect.

This study is the first to establish a correlation between a *KCNC1* variant and defects in membrane trafficking linked to disease. Using electrophysiological, biochemical, and imaging techniques, we demonstrated that the Ser474 residue is essential for the anterograde trafficking of the Kv3.1 channel. Previous research has exclusively described the trafficking mechanisms of Kv3.1 in neurons, which involve specific protein-protein interactions, such as the binding of KIF5B to the N-terminal of the channel for post-Golgi trafficking and the interaction of the C-terminal with Ankyrin-G for axonal entry (Barry et al., 2013; Xu et al., 2007; Xu et al., 2010). However, it remains unclear whether these proteins form a tripartite complex for axonal targeting or function in tandem trafficking. Both HEK293 and HeLa cells express transcripts for Ankyrin-G and KIF5B, suggesting a trafficking mechanism similar to that in neurons. Nonetheless, other molecular mechanisms for Kv3.1 trafficking cannot be excluded. End-binding (EB) proteins, which interact with microtubules and cargo proteins, may facilitate Kv3.1’s microtubule-dependent trafficking (Akhmanova & Steinmetz, 2008). Furthermore, EB1 and EB3 proteins bind to Ankyrin-G, promoting AIS stabilization (Leterrier et al., 2011), but the relevance of these interactions in non-neuronal models remains unclear.

Using molecular dynamics simulations, we analyzed the C-terminal folding of the wild-type and p.S474C variants of the Kv3.1b channel. Our results suggest that the p.S474C variant disrupts key interactions between residues P464, Y471, C472, V476 and S478 of one subunit and the amino acids B30, A35, W36 and E39 of the following subunit, which could be involved in channel folding. This disruption is evidenced by the altered folding dynamics observed in the simulations, where the S474C substitution exhibited instability and failed to reach a stable RMSD plateau, contrasting with the stable behavior of wild-type Kv3.1b. The structural changes induced by the S474C substitution could alter the channel’s functionality and membrane trafficking. Furthermore, in our model, the S474C substitution resulted in primarily maintaining interactions within the same subunit, as opposed to the inter-subunit interactions observed with the Kv3.1b^WT^. This shift in the interaction landscape is indicative of a broader destabilization that could compromise the channel’s functional efficacy. Although previous structural studies have shown the interactions between the residues D97, E98, D100 and E102 from the N-terminal T1 domain with the residues K454, K455, K456, K457 and K458 in the C-terminal region Kv3.1 modulate the channel gating and trafficking (Chi et al., 2022), this works used the Kv3.1a isoform for Cryo-EM studies which is 50 amino acids shorter than Kv3.1b explaining the differences with our modeling. Moreover, the proposed model for the AIS targeting of the Kv3.1b channel involves the exposure of the lysine cluster (K454-K458) leading to its interaction with Ankyrin G (Gu, 2012), supporting that the interaction between the T1 domain and the C-terminal of the Kv3.1b channel could depend on other surrounding residues (Kaczmarek, 2017). This supports our model for wild-type Kv3.1b, marking the first structural model of this ion channel, which could serve as a foundation for future structural studies.

Mutations in Kv channels affecting folding and assembly can lead to various inherited channelopathies. These disorders often involve altered neuronal excitability, cardiac arrhythmias, or muscle dysfunction (Harraz & Delpire, 2024; Waxman, 2001). For instance, *KCNQ2* and *KCNQ3* variants, encoding Kv7.2 and Kv7.3 channels, respectively, have been linked to impaired folding, trafficking, and membrane availability, resulting in decreased potassium conductance and neuronal hyperexcitability (Etxeberria et al., 2008; Liu & Devaux, 2014; Maljevic et al., 2011; Urrutia et al., 2021; Zhang et al., 2020). Another example is the Kv1.1 channel (*KCNA1*), which is associated with episodic ataxia type 1 (EA1) (Chen et al., 2016; Rea et al., 2002). Similarly, mutations in the Kv11.1 channel (hERG), encoded by the *KCNH2* gene, are implicated in Long QT Syndrome Type 2 (LQTS2), a cardiac disorder characterized by prolonged repolarization and an increased risk of arrhythmias (Nakano & Shimizu, 2016). Several variants in hERG channels have been shown to disrupt normal folding, leading to ER retention and degradation (Anderson et al., 2014; Hall et al., 2018).

Kv3.1b is expressed in high-frequency firing neurons, such as granule cells in the cerebellar cortex, neurons in the reticular nucleus of the thalamus, and parvalbumin containing (PV+) GABAergic interneurons in the hippocampus and neocortex (Kaczmarek & Zhang, 2017; Ozaita et al., 2002). The high activation threshold, fast activation, and rapid deactivation of Kv3.1b allow for quick repolarization during APs, shortening their duration (Kaczmarek & Zhang, 2017; Pongs, 2008; Rudy & McBain, 2001). Consequently, a loss of function in Kv3.1b in PV+ interneurons could reduce firing in these cells, leading to network hyperexcitability and contributing to various neurological disorders (Gunthorpe, 2022; Nascimento & Andrade, 2016).

Rett syndrome has been broadly linked to an imbalance in excitatory/inhibitory neurotransmission (Chao et al., 2010). Deleting *MECP2* in PV+ neurons in mouse models produces a Rett syndrome-like phenotype, including motor, sensory, memory, and social deficits (Ito-Ishida et al., 2015). Similarly, deleting Kv3.1 in mice results in social impairments (Bee et al., 2021), suggesting that dysfunction in PV+ interneurons may play a critical role in social behaviors. In this context, further research into the role of *KCNC1* in neurological diseases is necessary to understand how Kv3.1 loss of function can lead to diverse phenotypes in patients.

In summary, our findings identify and characterize a novel *KCNC1* variant in a patient with clinical manifestations consistent with Rett syndrome, which affects Kv3.1 channel trafficking and membrane availability, highlighting the importance of Kv3.1 trafficking in disease pathogenesis. Mutations that disrupt Kv3.1 trafficking can alter its expression at the cell membrane, affecting neuronal excitability and function. Several studies have explored the use of pharmacological modulators of Kv3.1 currents to influence neuronal excitability and treat various conditions, including hearing disorders, bipolar mania, ataxia, epilepsy, and other neurodevelopmental disorders (Ambrosino et al., 2023; Brown et al., 2016; Chen et al., 2023; El-Hassar et al., 2019; Parekh et al., 2018). Thus, understanding the mechanisms underlying Kv3.1 channel trafficking may provide therapeutic benefits for individuals with Kv3.1-associated conditions.

## Materials and Methods

### Study approval

Clinical case report was conducted according to Ethical Protocol (N° 170/2024) (Scientific Ethics Committee, Servicio Metropolitano de Salud, Chile). Animal experiments were conducted according to protocols approved by the Ethics Committee of the Universidad de Santiago de Chile (N° 301/2018), in accordance with the rules and guidelines of the National Research and Development Agency (ANID), Chile.

### DNA sequencing

Genomic DNA was isolated from peripheral blood using the *Wizard Genomic DNA Purification* Kit (Promega, Madison, USA), according to the manufacturer’s instructions. Primers flanking the *KCNC1*:c.1421C>G variant were designed using *PrimerQuest* (Integrated DNA Technologies, USA). PCR amplification of genomic DNA was performed using GoTaq Polymerase (Promega, USA) followed by conventional Sanger sequencing technology using a 3500xL Genetic Analyzer (Applied Biosystems) in “Unidad de Secuenciación y Tecnologías Ómicas de la Pontificia Universidad Católica de Chile”.

### *In silico* predictions

Prediction algorithms were run from the Varsome database (Kopanos, 2019. Available in https://varsome.com) based on *KCNC1* c.1421C>G (p.Ser474Cys) (ClinVar access: VCV000851860.8) and *RELN* c.7648A>G (p.Met2550Val) (ClinVar access: RCV001056357.7) using algorithms available in the dbNSFP 4.8 database (Liu et al., 2011; Liu et al., 2020). Meta algorithms used included BayesDel addAF and BayesDel noaddAF (Feng, 2017), MetaLR (Dong et al., 2015), MetaRNN (Li et al., 2022), MetaSVN (Dong et al., 2015) and REVEL (Ioannidis et al., 2016), using cutoff values suggested by Varsome. Population allele frequencies were obtained from the gnomAD 4.0 database (http://gnomad.broadinstitute.org).

### Cell culture and plasmids

HEK293 and HeLa cells were cultured at 37 °C, and 5% CO_2_ in DMEM High Glucose media (Invitrogen), supplemented with 5% v/v fetal bovine serum (FBS) and were transiently transfected with plasmids using lipofectamine 2000 (Invitrogen) as described. Plasmids encoding FLAG-Kv3.1^WT^ and FLAG-Kv3.1^S474C^ were obtained from Genescript (#NM_004976.4), Str-KDEL-IRES-mScarlet-i-Sec23a was obtained from Addgene (Addgene plasmid #117273). Kv3.1^WT^ and Kv3.1^S474C^ were subcloned in the mVenus-N1 vector (Addgene plasmid # 27793).

### Electrophysiological recordings

Whole-cell patch clamp experiments on HEK293 cells transiently transfected or in cortical neurons were performed using an intracellular solution containing (in mM) (Hideshima et al., 2001): 120 K-gluconate, 10 KCl, 8 NaCl, 0.5 EGTA, 4 Na-ATP, 0.3 Na-GTP, 10 HEPES, 10 Phosphocreatine, pH 7.2 adjusted with KOH, ∼300 mOsm/kg. The extracellular solution contained (in mM) (Hideshima et al., 2001): 140 NaCl, 2.5 KCl, 1 MgCl_2_, 2.5 CaCl_2_, 10 HEPES, and 10 glucose, adjusted to pH 7.4 with NaOH, 300 mOsm/kg. For voltage-clamp recordings, the voltage steps protocols consisted of a series of depolarizing voltage steps from −80 to 100 mV with a 10 mV increase with a duration of 0.5 s delivered somatically at 1 Hz from a holding potential of −80 mV, pipette and whole-cell capacitance was fully compensated and series resistance was compensated by 80%. For noise analyses, the voltage steps were applied at least 6 times each (at least 120 pulses per cell). The variance of the currents were calculated from the voltage steps, the data was plotted in an XY distribution and fitted to quadratic function 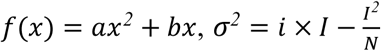to estimate the open probability, unitary conductance, and number of channels, where “*N*” correspond to number of channels, “*i*” correspond to unitary current and *I* correspond the macroscopic current, as described in (Alvarez et al., 2002). For current-clamp recordings, the protocol consisted in a series of depolarizing current steps from −60 to 160 pA with a 20 pA increase and a duration of 1 s, delivered at 1 Hz with a holding current of 0. Bridge balance and capacitance neutralization were applied. Liquid junction potential was 14.7 mV, and it was not subtracted (Marino et al., 2014) using LJPcalc software (https://swharden.com/LJPcalc). Electrophysiological recordings were performed using a Multiclamp 700A (Molecular Devices, San Jose, CA, USA) and digitized using a National Instruments PCIe-6323. Data was low pass filtered at 10 kHz and digitized at 50 kHz using WinWCP 5.7 (https://github.com/johndempster/WinWCPXE/releases/tag/V5.7.8). Data was analyzed using Clampfit 10.3 and Igor Pro 6.37 with the Neuromatic module (Rothman & Silver, 2018). Activation data were plotted as normalized conductance and were fitted to a single Boltzmann function.

### Immunoblot analysis

Cells were lysed in lysis buffer [50 mM Tris-HCl, 150 mM NaCl (Merck, Darmstadt, Germany, catalog #1064045000), 1 mM ethylenediaminetetraacetic acid (EDTA; Chemix, Lampa, Santiago, Chile, catalog #160408), 1 mM sodium orthovanadate (Calbiochem, San Diego, CA, USA, catalog #567540), 5 mM NaF (Sigma-Aldrich catalog #S7920), 1% v/v Triton X-100 (Sigma-Aldrich catalog #10789704001), pH 7.4] containing as protease inhibitors 1 mM phenylmethylsulfonyl fluoride (PMSF; Sigma-Aldrich catalog #78830) and a protease inhibitor cocktail (PIC; Cytoskeleton, Inc, Denver, CO, USA, catalog #PIC02) for 30 minutes at 4°C. The lysates were centrifuged at 11,000 g at 4°C for 10 minutes. Reducing Sample Buffer (RSB) [62.5 mM Tris-HCl, 2% w/v SDS (Sigma-Aldrich catalog #L5750), 10% v/v glycerol (Merck catalog #356352), 1% v/v β-Mercaptoethanol (Merck catalog #8057400250)] was added to the supernatant samples, and then boiled for 5 minutes followed by size fractionation on SDS–PAGE. Following SDS-PAGE, proteins were transferred to nitrocellulose membranes (GE Healthcare Life Sciences catalog #10600002), which were then blocked for 1 hour with BLOTTO [4% w/v nonfat dry milk/ 0.1% v/v Tween-20 in Tris-buffered saline (TBS: 50 mM Tris, pH 7.5, 150 mM NaCl)] followed by 2 hours or overnight incubation with primary antibodies. After 3 washes with BLOTTO, the membranes were incubated with the appropriate HRP-conjugated secondary antibody for 1 hour. After 3 washes for 10 minutes each with 0.1% v/v Tween-20/TBS, immunoblots were visualized by Pierce ECL Western Blotting Substrate (Thermo Fisher Scientific catalog #34080). The images were acquired with a Mini HD9 imager (Uvitec Ltd., Cambridge, UK) and quantified using the Nine-Alliance software (Uvitec Ltd.).

### Surface biotinylation assay

Biotinylation assays were performed as described (Cerda et al., 2015). The cells were grown to 80%-90% confluence on poly-L-lysine (200 µg/mL) treated 35 mm tissue culture dishes. The cells were washed twice with ice-cold DPBS (pH 8.0) and incubated with 1 mg/mL EZ-link sulfo-NHS-LC-biotin (Thermo Fisher Scientific catalog #21335) dissolved in DPBS (pH 8.0) for 30 minutes at 4°C. The reaction was terminated with a blocking solution (50 mM Tris, 154 mM NaCl, pH 8.0), and the cells were washed twice with ice-cold DPBS. The cells labeled with sulfo-NHS-biotin were lysed as described above, and lysates were incubated with streptavidin-agarose (Thermo Fisher Scientific catalog #20351) for 2 h at 4°C. The precipitated proteins were eluted with RSB and resolved in SDS-PAGE for further immunoblot analysis.

### Nocodazole synchronization

Synchronization of the trafficking based on Nocodazole was performed in HeLa cells transiently transfected with the FLAG-Kv3.1b or FLAG-Kv3.1b^S474C^ expressing for 48 h. The cells were incubated with 20 μM Nocodazole for 16 hours at 37°C in DMEM supplemented with 5% FBS (Storrie, 1998), then the drug was removed washing two times with DPBS and were incubated DMEM supplemented with 5% FBS at 37°C for 1 hour. Then, cells were fixed for further immunofluorescence assays.

### Immunofluorescence staining

Forty-eight hours post-transfection, cells were fixed for 15 minutes at 4°C in fixative solution [4% w/v formaldehyde (freshly prepared from paraformaldehyde, Sigma-Aldrich catalog #158127), 4% w/v sucrose (Sigma-Aldrich catalog #S0389) in Dulbecco’s phosphate buffered saline (DPBS), pH 7.4]. For surface staining experiments, cells were incubated with WGA-Alexa fluor 555 (Thermo Fisher Scientific catalog # W32464) in DPBS for 45 minutes at room temperature. Cells were then permeabilized and blocked with a blocking solution containing 0.1% v/v Triton X-100, 4% w/v nonfat dry milk in DPBS for 30 minutes at room temperature and incubated with the respective primary antibody (Table 1) for 1 hour at room temperature. After three washes for 10 minutes each, primary antibodies were detected by incubation for 1 hour at room temperature with Alexa-labeled isotype-specific secondary antibodies (Table 1) and Hoechst 33258 nuclear stain at 200 ng/mL (Thermo Fisher Scientific catalog #H3569). Samples were washed 3 times for 10 minutes each on 0.1% v/v Triton X-100/DPBS and mounted with Fluoromount (Sigma-Aldrich catalog #F4680). Images were acquired with 63× objective 1.35 N.A. using optical sectioning and structured illumination Zeiss Apotome 2, Axiovert 7 (Carl Zeiss, Oberkochen, Germany).

**Table 1:** List of antibodies used in this study.

| *Antibody* | Isotype | *Dilution* | *Final [],*<br>*μg/mL* | Source | Type | Catalog | RRID | Purification |
| --- | --- | --- | --- | --- | --- | --- | --- | --- |
| Anti-FLAG | IgG | 1:3000 (IB) | 0,4 | Sigma-Aldrich | pAb | F7425 | AB_439687 | AP |
| Anti-α-Tubulin | IgG1 | 1:5000 (IB)<br>1:2000 (IF) | 1<br>2,5 | Sigma-Aldrich | mAb | T5168 | AB_477579 | Asc |
| Anti-Golgin97 | IgG1 | 1:100 (IF) | 1 | Thermo Fisher Scientific | mAb | A-21270 | AB_221447 | - |
| Anti-Ankyrin-G | IgG2A | 1:100 (IF) | 1 | Addgene | mAb | 192039 | AB_1069771<br>8 | AP |
| Goat anti-Rabbit HRP-conjugated Antibody | IgG | 1:5000 (IB) | - | Jackson Immunoresearch laboratories inc. | pAb | 11103514<br>4 | AB_2307391 | - |
| Goat anti-Mouse HRP-conjugated Antibody | IgG | 1:5000 (IB) | - | Jackson Immunoresearch laboratories inc. | pAb | 11503500<br>3 | AB_1001528<br>9 | - |
| Goat anti-Rabbit (H+L) Alexa fluor 488-conjugated antibody | IgG | 1:1000 (IF) | 2 | Thermo Fisher Scientific | pAb | A-11008 | AB_143165 | AP |
| Goat anti-Mouse IgG2A Alexa fluor 555-conjugated antibody | IgG | 1:1000 (IF) | 2 | Thermo Fisher Scientific | pAb | A-21137 | AB_2535776 | AP |
| Goat anti-Mouse IgG1 Alexa fluor 647-conjugated antibody | IgG | 1:1000 (IF) | 2 | Thermo Fisher Scientific | pAb | A-21240 | AB_2535809 | AP |

AP, affinity purified; Asc, purified from mouse ascites fluid by affinity chromatography; RRID, Research Resource Identifier.

### Retention Using Selective Hooks assay

Synchronization of the trafficking based on the Retention Using Selective Hooks system was performed in HeLa cells transiently transfected with the Kv3.1b^WT^-mVenus-SBP or Kv3.1b^S474C^-mVenus-SBP expressing plasmid, ERStreptavidin Hook and ERES marker (Str-KDEL-IRES-mScarlet-i-Sec23a) for 48 h. Kv3.1b-mVenus-SBP trafficking was induced by addition of 40 mM final concentration of D-biotin for 0, 30, 60, 120 and 180 min, as previously described (Blanco et al., 2019; Boncompain et al., 2012; McCaughey et al., 2019). Then, cells were fixed for further immunofluorescence assays.

### Kv3.1b channel modeling and molecular dynamics simulation

The cryo-EM Kv3.1 channel structure (PDB ID: 7PQT) was used as a template. The missing C-terminal segment in the crystallographic structure (residues 454-585) was generated using CollabFold and overlapped with PyMOL to the templated structure. The generated model was refined using the coarse-grained refinement provided by HADDOCK, and finally, PRAS-SERVER was used in the refined structure to avoid nomenclature problems by adding missing hydrogens and fixing the side chains. PyMOL was used to generate the S474C mutation in the model. The Kv3.1 wild-type and Kv3.1^S474C^ systems were prepared for molecular simulation. The Kv3.1b channel was embedded in a palmitoyl oleoyl phosphatidylcholine (POPC) lipid membrane, itself contained in a box with pre-equilibrated TIP3P-type waters under periodic boundary conditions. The system was neutralized and a concentration of 0.15 NaCl was added through the Autoionize plugin of the Visual Molecular Dynamics (VMD) program. The CHARMM36 force field under isobaric-isothermal (NPT) conditions was used to run molecular simulations. A molecular dynamics simulation (MDs) of the previously constructed system was launched through the high-performance molecular simulation software NAMD. The following minimization and equilibration protocol was used: (I) Minimization by 30,000 steps to decrease and harmonize steric interactions in the system (II) Lipid heating by 0.5 ns, (III) Relaxation of membrane and solvent with constrained proteins (0.5 ns), (3) Relaxation of side chains where the protein backbone was constrained until it was completely free (1 ns), and (4) Relaxation of the whole system (1∼250 ns). Finally, 500 ns of production of molecular simulation. RMSD calculations were performed in Visual Molecular Dynamics (VMD). The intermolecular interactions formed by residue 474 along the molecular dynamics trajectory of the wildtype and S474C systems were calculated using GetContacts. A cutoff of 10 Å along the molecular dynamics was used to measure the residues that maintains close contacts with residue 474 in the WT and S474C systems through a TCL script executed in VMD. From the results of the C-terminal folding mutation effect of the Kv3.1b model, the chain of the structure with the stable RMSD of the trajectory (chain A) was extracted and used to tetramerize the Kv3.1b channel-based on this monomer. The resulting Kv3.1b channel model was used to evaluate the effect of the S474C mutation on DUET (Pires, 2014), DDMut (Rodrigues, 2021), DynaMut2 (Rodrigues, 2021), and mCSM (Pires, 2014) web servers. These prediction servers estimate the effect of single-point mutations based on calculations of unfolding free energy (ΔΔG_Mut→WT_) defined as:

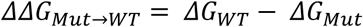

The predicted values are expressed as the variation of Gibbs free energy (kcal/mol) where positive values correspond to stabilizing mutations and negative values denote destabilizing mutations.

### Cortical neuronal cultures

Primary frontal cortical neurons were prepared from E18 C57BL/6J mouse embryos. The frontal cortices were dissected in Hank’s balanced salt solution (HBSS) and digested in trypsin (0.25 % w/v) plus DNase I (0.03 mg/mL) for 8 minutes at 37°C, triturated, and plated in Minimum essential media (MEM) supplemented with horse serum (20%v/v), glucose (0.1% w/v), sodium pyruvate (0.5mM), HEPES (10 mM), and penicillin-streptomycin (100 I.U./mL). Neurons were plated at a density of 50,000 cells per well on 12 mm coverslips pre-coated with poly-D-lysine (30 µg/mL) and laminin (2 µg/mL). After 6 hours of plating, the media was replaced with Neurobasal medium supplemented with 2% B27, GlutaMAX-I (1%), and penicillin-streptomycin (100 I.U/mL). Neurons were cultured at 37°C with 5% CO_2_. Half of the culture media was replaced every 3 days. Experiments were performed between days *in vitro* (DIV) 10-12.

### Simulation of neuronal excitability

Following (Gu et al., 2012), and with minor modifications, we adopted a simplified geometry with three cylindrical compartments: dendrite, soma and axon. Dendrite was 250 µm long and with a diameter starting at 3 µm next to the soma, increasing to 5 µm at 60 µm distance, and decreasing to 4 µm at the farther end. Soma was 20 µm long and 20 µm in diameter. Finally, the axon had a length of 800 µm and a diameter of 1 µm. The dendrite was spatially discretized into 41 segments, the axon in 201 segments, and the soma was considered as a single iso-potential compartment. They were connected with an internal specific resistivity R_a_ of 200 Ωcm.

In each segment, the dynamics of membrane voltage is given by the current balance equation:

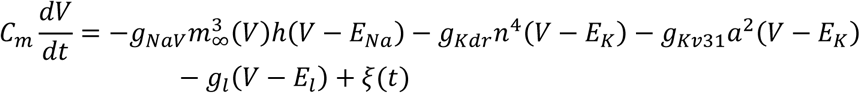

where g_NaV_, g_Kdr_, and g_Kv3.1_ represent the maximum conductances of the voltage-dependent Sodium (NaV) channels, generic delayed-rectifier Potassium (Kdr) channels and the Kv3.1b channels, respectively. gl is the voltage-independent leak conductance. E_Na_, E_K_, and E_l_ are reversal potentials for Sodium, Potassium, and leak currents, respectively. m_∞_ is the activation gate of NaV channels, considered here to be instantaneous.

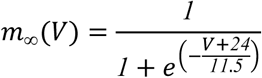

*h*, *n*, and *a* are gating variables that obey the differential equation

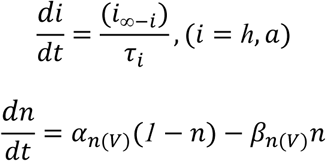

Where

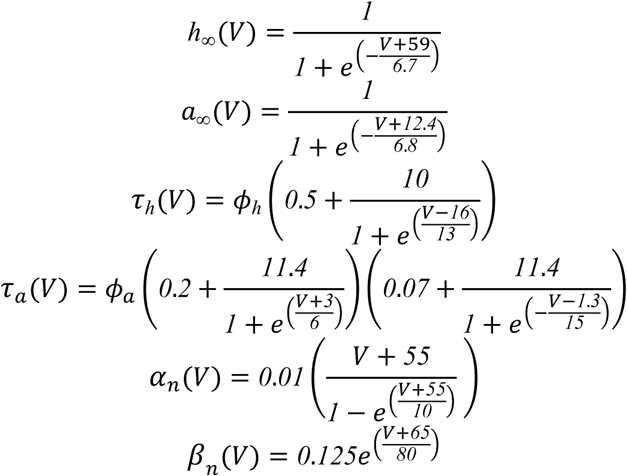

ξ(t) is a noise current term implemented as an Ornstein-Uhlenbeck process, such that *dξ*/*dt* = (*χ*(*t*) − *ξ*)/*τ*_*ξ*_ with τ_ξ_=3 ms. χ(t) is a normally distributed random variable with 0 mean and standard deviation equal to 0.01 mA/cm^2^.

Other parameters are as follow: ϕ*_h_*=0.8; ϕ*_a_*=0.2, E_Na_=50, E_K_=-90, E_l_=-70 mV. In the soma and dendritic compartment, g_NaV_=0.1865; g_Kdr_=0.005; g_Kv3.1_=0.2; g_l_=8×10^-5^ mS/cm^2^. In the axon, g_Kdr_=0.004; g_Kv3.1_=0.8; g_l_=8×10^-5^ mS/cm^2^. Finally, the axonic value of g_NaV_ was 2.5 mS/cm^2^ in the segment between 10 and 45 µm from the soma (axon initial segment) and 0.746 mS/cm^2^ elsewhere. The model was implemented and run in NEURON 8.2.6 using Python 3.12 scripts (Hines et al., 2009)

### Statistical analyses

The data shown correspond to the mean ± SEM of at least three independent experiments. The data were analyzed using a two-tailed Mann–Whitney test to compare two conditions. For multiple comparisons, one-way analysis of variance (ANOVA) tests and Dunnett’s post-test were applied. For I/V curves, two-way analysis of variance (ANOVA) tests and Tukey’s post hoc tests were applied. Analyses were performed using GraphPad Prism v8.0 (GraphPad Prism, San Diego, CA, USA).

## Author contributions

DM performed research, analyzed data, and contributed to manuscript writing. CR collected and analyzed data and contributed to manuscript writing. JL analyzed data and contributed to manuscript writing. PS collected data. HM analyzed data. IS performed research and analyzed data. PC performed research and analyzed data. DR performed research. JB performed research and analyzed data. WG analyzed data. PO performed research and analyzed data. ES contributed to manuscript writing. MT collected and analyzed data. ELS designed the research, analyzed data, and contributed to manuscript writing. OC conceived the study, designed the research, analyzed data, and contributed to manuscript writing.

## Data Availability Statement

Data from this paper is available from the corresponding author upon request.

## Acknowledgments

We thank Mr. Nicanor Villarroel, Ms. Ana María Ramírez and Mr. Rodrigo Santos for their technical support. This research was funded by FONDECYT Grant 1240633 (to OC), the Millennium Nucleus of Ion Channel-Associated Diseases (to OC and WG), FONDECYT Grant 1220680 (to ELS) and Proyecto Postdoc DICYT, codigo 022243LS_Postdoc, Vicerrectoria de Investigacion, Desarrollo e Innovación (to ELS). FONDECYT Grant 1230446 funds WG. FONDECYT Grant 1241469 and Centro Basal FB0008 fund to PO. DM, JL, and JB are recipients of Beca Doctorado Nacional (#21201941, #21230481 and #21210923, respectively). ANID Postdoctoral Fellowship 3240187 funds to ES.

## Figure Legends

**Supplementary Figure 1.**
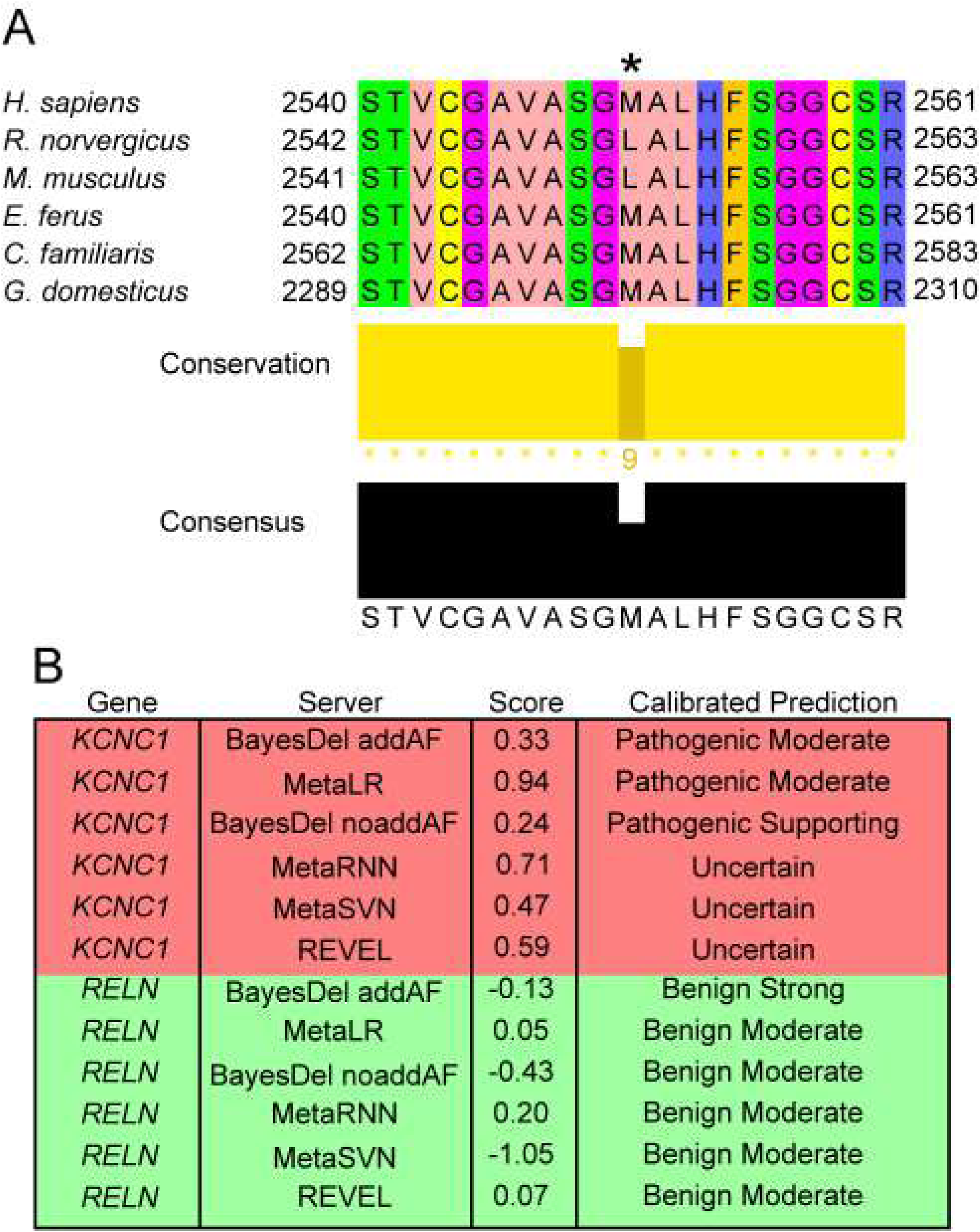
p.M2550T in *RELN* is benign by computational analysis. **(A)** Sequence alignment of Reelin protein sequences from *Homo sapiens* (human), *Rattus norvergicus* (rat), *Mus musculus* (mouse), *Equus ferus* (horse), *Canis familiaris* (dog), and *Gallus domesticus* (chicken). A multiple-sequence alignment was built with ClustalW server and analyzed with Jalview v.1.8.3. The sequence alignment is colored according to the chemical properties of the residues. Conservation of the S474 residue is indicated (asterisk). Conservation and consensus are indicated in the bottom diagrams. M2551 residue is labeled with asterisk. **(B)** Calibrated pathogenic prediction for KCNC1 and RELN genes obtained from different *in silico* predictors.

**Supplementary Figure 2.**
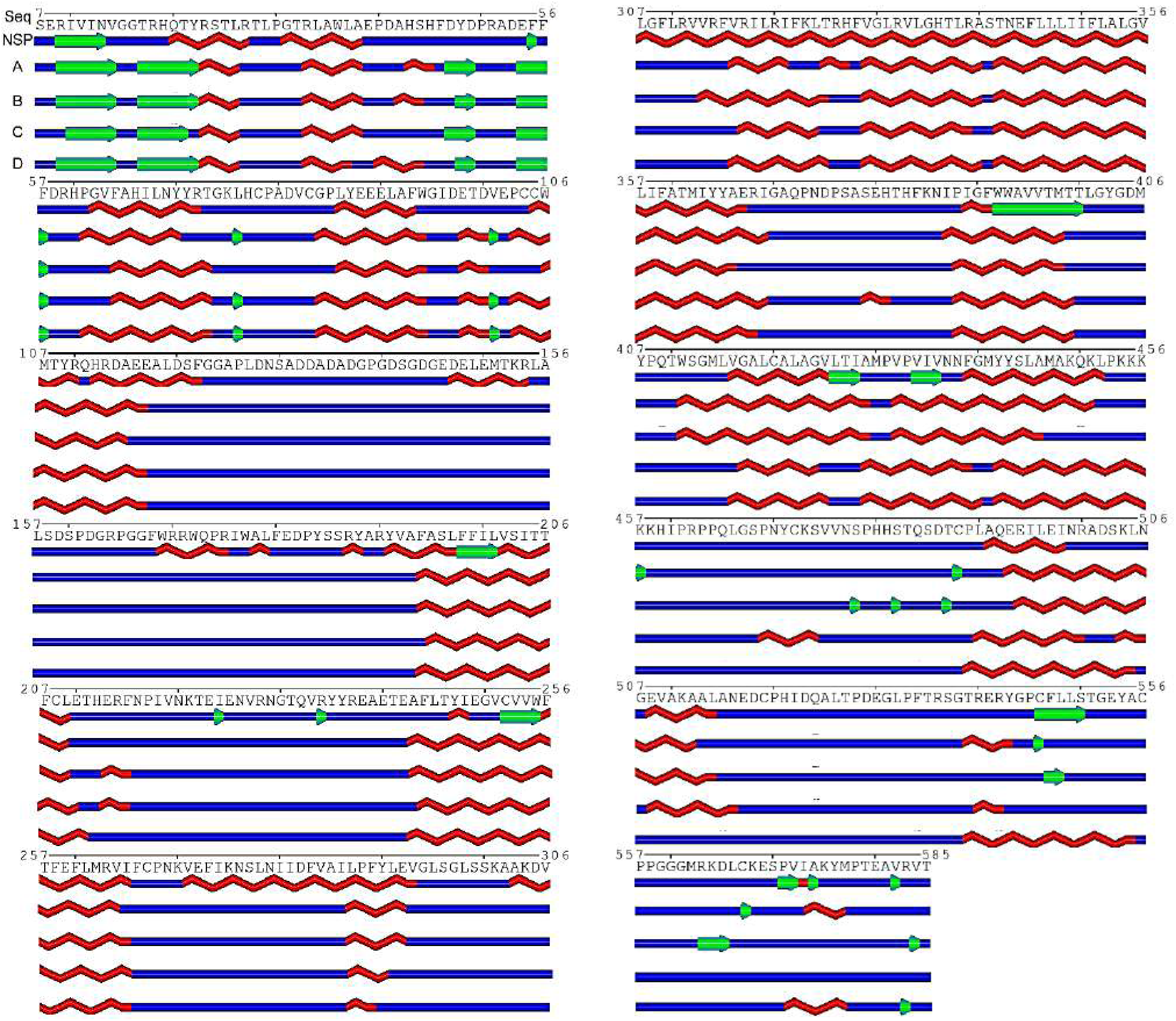
Secondary structure prediction analysis. Comparison of the consensus secondary structure prediction made in NSP with the secondary structure corresponding to the last molecular dynamics frame of each chain. α-helices, β-sheets and loops are shown in red, green and blue, respectively. Polyview was used to generate the scheme.

